# Biological modifications of the immune response to COVID-19 vaccine in patients treated with rituximab and immune-checkpoint inhibitors

**DOI:** 10.1101/2024.03.05.583494

**Authors:** Francesco Ravera, Martina Dameri, Isabella Lombardo, Mario Stabile, Neri Fallani, Camilla Scarsi, Benedetta Cigolini, Giusy Gentilcore, Alexander Domnich, Lodovica Zullo, Eugenia Cella, Giulia Francia, Eugenia Montanari, Andrea Orsi, Andrea Bellodi, Fabio Ferrando, Darawan Rinchai, Filippo Ballerini, Bianca Bruzzone, Damien Chaussabel, Jean-Charles Grivel, Carlo Genova, Roberto Massimo Lemoli, Davide Bedognetti, Alberto Ballestrero, Lorenzo Ferrando, Gabriele Zoppoli

**Affiliations:** Department of Internal Medicine and Medical Specialties, University of Genoa, Genova, Italia; IRCCS Ospedale Policlinico San Martino, Genova, Italia; Sidra Medicine, Doha, Qatar; Department of Experimental Medicine, University of Genoa, Genova (Italia); Department of Infectious diseases, St. Jude Children Research Hospital, Memphis, TN, USA; The Jackson Laboratory, Bar Harbor, ME, USA

**Keywords:** Rituximab, Immune-checkpoint inhibitors, COVID-19 vaccine, RNA sequencing, Interferon

## Abstract

Investigating the impact of immune-modulating therapies on mRNA vaccine efficacy transcends the immediate context of the COVID-19 pandemic. This study focuses on the differential immune responses to the third dose of COVID-19 mRNA vaccine among healthy volunteers, cancer patients treated with immune-checkpoint inhibitors (ICIs), and those treated with the anti-CD20 antibody rituximab. Utilizing RNA sequencing, serology, and interferon-γ release assessment, we charted the temporal dynamics of the immune response in such cohorts. Our findings indicate that ICIs maintain an immune profile similar to that of healthy individuals, whereas treatment with rituximab is associated with impairment of type I interferon response and the upregulation of transcripts pertaining to regulatory T cells, with a global dysfunction of both humoral and cellular immunity. This research deepens our understanding of the sophisticated interplay within the immune system in health and disease states, potentially informing therapeutic strategies across a spectrum of immunological conditions.

**Significance statement:** Our study examines how cancer treatments that modify the immune system affect transcriptional, serological, and cellular responses to a model for repeated antigenic stimulation in humans, represented by the SARS-CoV-2 booster vaccine. Specifically, we investigated patients treated with rituximab (RTX), which impairs antibody production, and immune checkpoint inhibitors (ICI), which can trigger autoimmune disorders. We discovered that RTX-treated patients not only exhibit a reduced antibody response but actually show a diminished interferon-mediated immune response, indicating a broader immune disruption than anticipated. Conversely, ICI-treated patients responded to the vaccine similarly to healthy individuals, suggesting that fears of adverse vaccine reactions in these patients may be unfounded. This research highlights important considerations for the clinical management of cancer patients receiving these treatments.

## Introduction

The advent of mRNA vaccines has represented an instrumental game-changer in the fight against the Coronavirus disease 2019 (COVID-19) pandemic. Specifically, the primary vaccination cycle with BNT162b2 or mRNA-1273 offered a 95% protection against the original SARS-CoV-2 strain in the general population [1,2]. However, this protection wanes after six months, making a third (booster) dose necessary [3]. Cancer patients, being especially at risk, have prompted extensive research into their vaccine-induced immune response [4,5].

Immune perturbagens commonly used in cancer treatment, such as anti-CD20 agents and immune-checkpoint inhibitors (ICIs), exert distinct immunosuppressive and immunostimulatory effects on the host immune system [6,7]. Among cancer patients, those with hematological malignancies treated with anti-CD20 agents show the lowest seroconversion rate to COVID-19 vaccination [4,8]. In contrast, treatment with ICIs has been linked with higher seroconversion rates compared to standard cytotoxic chemotherapy, suggesting a potential enhancing effect of immunotherapy on vaccination [9].

Vaccination represents an ideal model for the evaluation of the response mechanisms to a reproducible antigenic stimulation. Indeed, systems vaccinology has shed light on the intricate interplay between innate and adaptive immunity in the physiologic response to mRNA vaccines, revealing distinct response patterns after the first and second dose [10,11]. Notably, Arunachalam *et al.* identified distinct early interferon responses following the first and second dose of the BNT162b2 vaccine, with an enhanced innate immune response, particularly after the second one [10]. This heightened response is characterized by an increase in CD14^+^ CD16^+^ monocytes and antigen presenting cells, which facilitate the crosstalk between B and T cells.

A study by Lee *et al.* on patients with chronic lymphocytic leukemia, using blood RNA-Sequencing (RNA-Seq), revealed an early interferon response akin to that of cancer-free individuals [12]. However, limitations such as low sample size, absence of cancer-free controls, and heterogeneity of treatments and vaccination strategies precluded definitive conclusions.

Overall, the effects of iatrogenic perturbation of the adaptive immunity, whether through B-cell depletion or T-cell stimulation, on the immune response to COVID-19 vaccination in patients with blood and solid cancer, remain largely underexplored.

In this study, we conducted a comprehensive, serial analysis of the immune response to the booster dose of COVID-19 mRNA vaccines by contrasting its effects in healthy individuals and in cancer patients undergoing immune-system-perturbing therapies. In particular, we compared cancer-free volunteers (C-FV) with patients affected by non-Hodgkin lymphoma treated with anti-CD20 agents (rituximab - RTX) and solid tumor patients on ICIs. We explored the evolving phases of vaccine-induced immune response utilizing whole blood RNA-Seq, complemented with multiplex serology and interferon-γ release assessment (IGRA) to evaluate the effectors of humoral and cell-mediated immunity, respectively. Building on our previous research [11], which dissected the transcriptomic temporal dynamics of the immune response to the primary mRNA vaccine cycle, we focused our RNA-Seq analysis on the early peak responses previously observed after the first and second dose (i.e. 24 hours and 5 days after each dose). Additionally, we broadened our analysis to include assessments of the humoral and cell-mediated immunity at six months post-booster, aiming to evaluate the enduring nature of the vaccine-induced immunity.

In summary, this study provides a comprehensive analysis of the immune response to the COVID-19 mRNA booster vaccine in cancer-free individuals, as well as cancer patients undergoing immune perturbagens such as anti-CD20 agents and ICIs. This analysis aims to delineate the specific impacts of different immune-perturbing treatments on the integrated dynamics and persistence of vaccine-induced immunity, expanding our understanding of immune responses in these distinct patient groups.

## Results

### Demographics and study design

We collected 214 peripheral blood samples from a total of 44 individuals over an eight-month time span, comprising 22 C-FV, 11 ICI patients, and 11 RTX patients. Clinical characteristics of the study cohorts are reported in **Table 1**, whereas collection and event timelines are depicted in **Supplementary Figure 1**.

**Table 1.** Study cohorts’ characteristics.

|  | C-FV<br>(N = 22) | RTX<br>(N = 11) | ICI<br>(N = 11) |
| --- | --- | --- | --- |
| **Age (IQR)** | 41 (36 - 56) | 65 (58 - 77) | 62 (57 - 76) |
| **Sex** |  |  |  |
| *Male* | 7 (32%) | 6 (55%) | 4 (36%) |
| *Female* | 15 (68%) | 5 (45%) | 7 (64%) |
| **Stage** |  |  |  |
| *Early* | - | 7 (64%) | 1 (9%) |
| *Advanced* | - | 4 (36%) | 9 (82%) |
| *NA* | - | - | 1 (9%) |
| **COVID-19** |  |  |  |
| *COVID-19*<br>*diagnoses* | 9 (41%) | 3 (27%) | 1 (9%) |
| *Pre-booster dose* | 2 (9%) | - | 1 (9%) |
| *Post-booster* | 7 (32%) | 3 (27%) | - |
| **Sampled**<br>**participants** |  |  |  |
| *Day 0* | 22 | 11 | 11 |
| *Day 1* | 22 | 11 | 11 |
| *Day 5* | 22 | 10 | 11 |
| *Week 4* | 22 | 9 | 11 |
| *Month 6* | 20 | 9 | 10 |
| **Median temporal interval**<br>**Day 0 - Week 4**<br>**(IQR)** | 27 (26 - 28) | 26 (26 - 27) | 26 (26 - 28) |
| **Median temporal interval**<br>**Day 0 - Month 6**<br>**(IQR)** | 194 (188.5 - 203) | 186 (175.5 - 194.5) | 196 (174 - 199.5) |

All RTX patients were diagnosed with non-Hodgkin lymphoma and were receiving rituximab at the time of enrollment, either as monotherapy (n = 9) or in combination regimens (bendamustine, n = 1 and R-COMP, n =1). The ICI cohort included patients with lung cancer (n = 8), liver cancer (n = 1), urothelial cancer (n = 1), and breast cancer (n = 1), treated with pembrolizumab (n = 9) or atezolizumab (n = 2). The majority of participants (n = 41) received the BNT162 vaccine booster, while three C-FV individuals were administered the mRNA-1273 vaccine.

The C-FV cohort was characterized by a significantly lower median age compared to both ICI patients (adjusted p-value = 0.0036) and RTX patients (adjusted p-value = 0.0021). Extended data, including tumor type and stage, comprehensive therapy details, comorbidities, COVID-19 vaccination history and diagnosis, and sample collection timelines, are provided in **Supplementary Table 1**.

To accurately profile the vaccination response in time, while providing minimal discomfort to the study participants, we utilized an ultra-low volume sampling procedure via fingerpick for both blood RNA-Seq and serological analyses, as previously described [11]. Blood transcriptome was assessed for all participants at baseline (day 0), 24 hours (day 1), and five days (day 5) following the booster dose. Serological profiling was performed at day 0, four weeks (week 4), and six months (month 6) post-booster. For SARS-CoV-2, we tested the reactivity to a stabilized trimer of the Spike protein (S-Trimer), its Receptor Binding Domain (RBD), the Spike subunit S1 (S1), and the Nucleocapsid (Nucleo) and Envelope (Env) proteins. Reactivity to the S1 antigens of various human coronaviruses (hCoVs), including SARS-CoV, MERS, HCoV-229E, HCoV-NL63, HCoV-HUK1, and HCoV-OC43, was also tested. The seroreactivity to these antigens was quantified by measuring the Binding Antigen Index (BAI) increment for IgM, total IgG, and the different subclasses of IgG (IgG1, IgG2, IgG3), as well as IgA (IgA1, IgA2). The cell-mediated immune response to the vaccine was assessed using IGRA in a subset of 12 C-FV, 4 ICI, and 8 RTX at day 0, day 5, week 4, and month 6. An interferon-γ concentration of 200 mUI/mL after stimulation with native SARS-CoV-2 S1 was set as the threshold for positive response, while lower values were considered as borderline or negative. A detailed description of the analyses is provided in the Methods.

### Serological profiling of C-FV identifies distinct patterns of humoral response to SARS-CoV-2 at four week and six months from vaccination

We assessed the humoral response to the COVID-19 booster dose by performing a comprehensive evaluation of antibodies directed against various antigens of SARS-CoV-2 and the S1 of several hCoVs. To evaluate vaccination response, we used a mixed-effects model, without and with intervening COVID-19 diagnosis as a covariate adjustment. This allowed us to quantify the variation of BAI to reflect both temporal antibody level changes (BAI*_t_*) and the association of such variation with COVID-19 diagnosis (BAI_COVID-19_).

In the C-FV group, the COVID-19 booster administration resulted in a notable increase in vaccine-targeted antibodies at week 4 (**Figure 1**), particularly total IgG targeting RBD (BAI*_t_* = 8.92, false discovery rate (FDR) = 1.17 10^-6^), S-Trimer (BAI*_t_* = 14.06, FDR = 7.62 10^-7^), and S1 (BAI*_t_* = 20.39, FDR = 7.62 10^-7^). No significant response to Nucleo was detected, except in one individual with prior virus exposure who showed a marked increase in anti-Nucleo total IgG and IgG2 levels (**Figure 1A**). A moderate increment in anti-Env total IgG was noted (BAI*_t_* = 0.022, FDR = 1.4 10^-4^). At month 6, a significant rise in antibodies associated with natural infection was detected, such as anti-Env IgM (BAI*_t_* = 0.037, FDR = 0.005) and total IgG (BAI*_t_* = 0.23, FDR = 0.00025), as well as anti-Nucleo IgM (BAI*_t_* = 0.049, FDR = 0.00062), regardless of COVID-19 diagnosis. Most vaccine-targeted antibodies showed further increases compared to week 4 (**Supplementary Table 2**).

**Figure 1.**
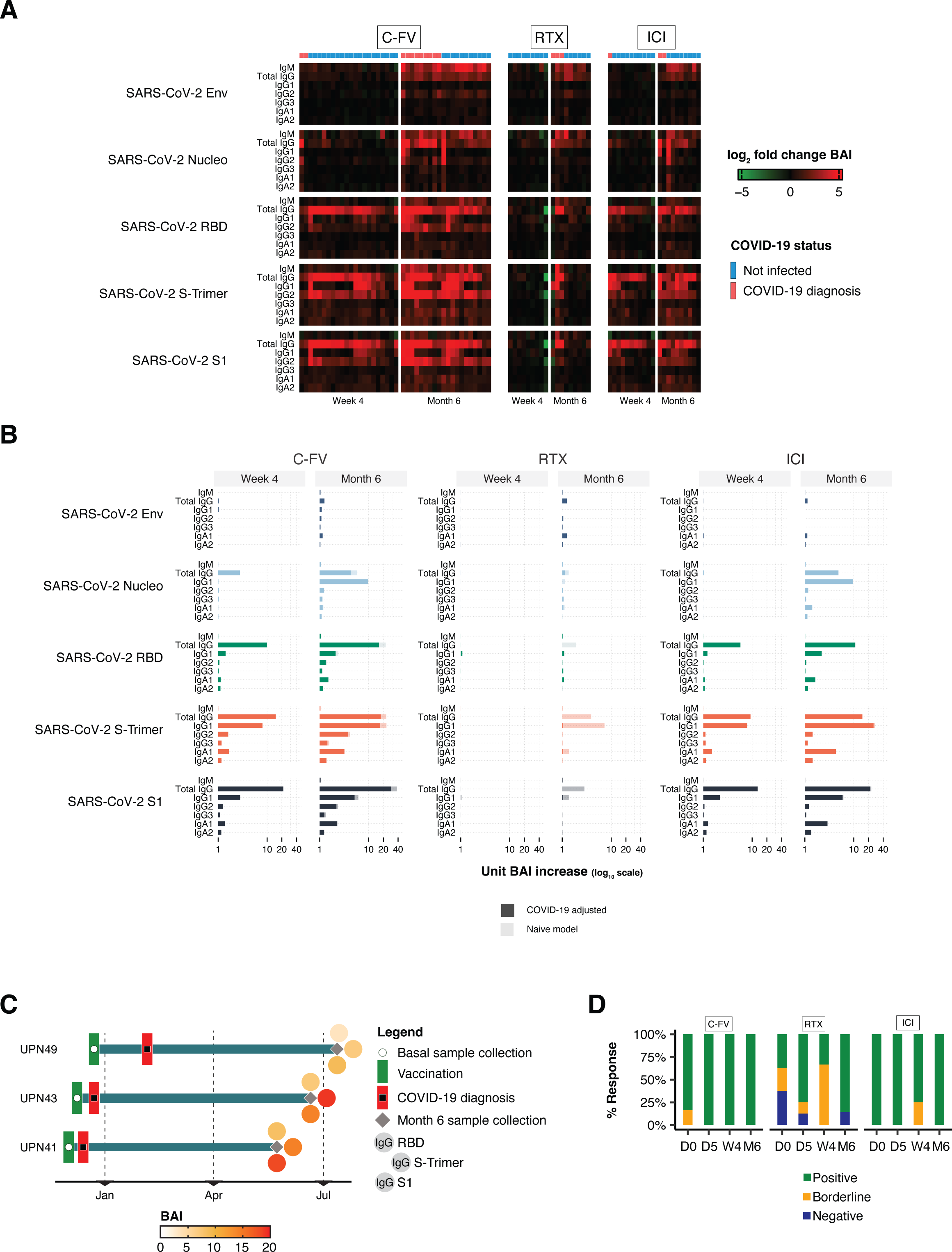
Humoral and cell-mediated immune responses after the third dose of the COVID-19 vaccine in C-FV, RTX, and ICI cohorts. A) Heatmap illustrating fold changes in Binding Antibody Index (BAI) for immunoglobulins targeting SARS-CoV-2 antigens: Envelope (Env), Nucleoprotein (Nucleo), Receptor Binding Domain (RBD), Full Spike Trimer (S-Trimer), and Spike S1 (S1). Red signifies an increase in BAI relative to baseline, while green denotes a decrease. Rows display immunoglobulin changes for IgM, total IgG, IgG subclasses (IgG1, IgG2, IgG3), and IgA subclasses (IgA1, IgA2), across time points (week 4 and month 6 versus baseline). The top annotation bar shows cumulative COVID-19 diagnosis status. B) Bar plots depicting unitary BAI changes at week 4 and month 6 versus baseline, with solid colors indicating adjustments for COVID-19 diagnosis in a mixed-effects model, and transparent colors for unadjusted estimates. C) Timeline highlighting the interval between vaccination and COVID-19 diagnosis in RTX patients, with month 6 total IgG-targeted BAI. D) Bar plots represent IGRA test results as percentages within each cohort, color-coded for positive (green), borderline (yellow), and negative (blue) outcomes at each time point.

Considering hCoVs, a pronounced cross-reaction with the SARS-CoV Spike protein was observed following the booster administration, as evidenced by a substantial augmentation in anti-S1 total IgG for SARS-CoV at week 4 (BAI*_t_* = 22.47, FDR = 4.32 10^-5^; **Supplementary Figure 2**), corroborating prior findings [11]. At month 6, there was a significant enhancement in anti-S1 IgM and total IgG for MERS (FDR of 0.0097 and 0.0073), HCoV-229E (FDR of 0.0031 and 3.34 10^-7^), HCoV-NL63 (FDR of 0.01383 and 0.0009), HCoV-HUK1 (FDR of 0.0092 and 0.0006), and HCoV-OC43 (FDR of 0.015 and 0.0012; **Supplementary Figure 2**; **Supplementary Table 3**). Anti-S1 IgM of hCoVs assessed at month 6 significantly correlated with anti-Env and anti-Nucleo IgM of SARS-CoV-2, independently of intercurrent COVID-19 diagnosis (**Supplementary Figure 3**). A similar trend was noted for SARS-CoV-2 anti-RBD IgM. Notably, SARS-CoV-2 anti-S-Trimer IgM correlated with anti-S1 IgM of hCoVs only in subjects without a diagnosis of COVID-19, whereas SARS-CoV-2 anti-S1 IgM showed correlation specifically in infected subjects. Taken together, our data suggest that multiple factors contribute to the development of a comprehensive and long-lasting humoral immunity to SARS-CoV-2, with infection by SARS-CoV-2 or other hCoVs possibly acting as a natural booster in previously vaccinated subjects.

### Humoral immune response to COVID-19 booster is preserved in ICI patients, but severely impaired in RTX cases

In ICI patients, the humoral response to the COVID-19 booster dose mirrored that of the C-FV group, exhibiting a pronounced increase in vaccine-specific antibodies at week 4. This included a rise in total IgG targeting RBD (BAI*_t_* = 4.75, FDR = 0.038, S-Trimer (BAI*_t_* = 8.19, FDR = 0.038), and S1 (BAI*_t_* = 11.90, FDR = 0.038 3.8 10^-2^). Similarly to C-FV individuals, ICI patients demonstrated a significant increase in infection-associated antibodies at month 6, including anti-Env IgM (BAI*_t_*= 0.016, FDR = 0.038) and total IgG (BAI*_t_* = 0.13, FDR = 0.013) and anti-Nucleo IgM (BAI*_t_* = 0.039, FDR = 0.089; **Supplementary Table 2**).

In contrast, RTX patients did not mount a humoral response to the booster dose, as shown by negligible changes compared to baseline in anti-RBD total IgG (BAI*_t_*= - 1.21, FDR = 0.70), anti-S-Trimer total IgG (BAI*_t_* = −1.60, FDR = 0.70), and anti-S1 total IgG (BAI*_t_* = −2.61, FDR = 0.70) at week 4. However, at month 6, we observed an increase in vaccine-targeted antibodies associated with intercurrent clinically manifest COVID-19 (anti-RBD total IgG BAI_COVID-19_ = 5.77, FDR = 0.074; anti-S-Trimer total IgG BAI_COVID-19_ = 13.08, FDR = 0.017; anti-S1 total IgG BAI_COVID-19_ = 13.32, FDR = 0.062; **Supplementary Table 1**). Indeed, three RTX patients were diagnosed with COVID-19 during the study period. Intriguingly, only patients UPN41 and UPN43, who contracted COVID-19 shortly after vaccination (12 and 14 days, respectively), mimicking the temporal proximity of the two vaccine doses of primacy vaccination cycle, developed a humoral response towards SARS-CoV-2 (**Figure 1C**). In contrast, patient UPN49, diagnosed with COVID-19 at 44 days post-booster, showed no response at month 6. While anecdotal, this finding suggests that a complete two-dose re-vaccination cycle may achieve higher seroconversion rates than a single booster dose in this subset of immunocompromised patients.

### The cell-mediated immunity against SARS-CoV-2 is impaired in RTX patients and is temporarily restored by the booster dose of COVID-19 vaccine

We evaluated the cell-mediated immune response to the booster dose using an IGRA assay measuring interferon-γ release from T-cells upon exposure to SARS-CoV-2 S1. C-FV group and patients in the ICI cohort showed no significant differences, all maintaining positive baseline IGRA results across all post-booster time points (**Figure 1D, Supplementary Table 4**).

Conversely, RTX patients displayed weaker cell-mediated responses than C-FV at day 0 and at week 4, with only 37.5% and 33.3% testing IGRA-positive, compared to 83.3% and 100% in C-FV (p-values of 0.062 and 0.0049, respectively). Of note, the cell-mediated response in such cohort was restored at day 5 and at month 6 after vaccination with 75% and 85.7% showing positive IGRA results.

This pattern in RTX patients suggests that prolonged rituximab exposure adversely affects cell-mediated immunity to SARS-CoV-2, likely due to disrupted B and T-cell interaction or, potentially, direct effects of rituximab on T cells [7]. Although the booster temporarily restores cell-mediated immunity, a persistent depletion of CD20+ cells may hinder, at least partially, a durable and efficient T cell-mediated response. The eventual recovery of this response at month 6, possibly influenced by infection by SARS-CoV-2 or other hCoVs (as noted for humoral responses), suggests reversibility and potential enhancement of cell-mediated immunity through repeated antigen exposure even in RTX patients.

### Modular analysis of COVID-19 booster reveals orchestrated immune response with early interferon spike followed by activation of adaptive immunity

To investigate the early dynamics of the immune response to the COVID-19 booster vaccination, we conducted a time-course analysis of the whole peripheral blood transcriptome through RNA-seq, applying the modular approach originally established by Chaussabel *et al.* and implemented in the BloodGen3 package [13–15]. This method involves categorizing RNA transcripts into 382 fixed transcriptional modules, co-clustering across 16 different immunological and physiological states and further consolidated in 38 aggregates for functional annotation. This framework enables the analysis of transcriptional module profiles at both group- and single-sample level, facilitating the identification of phenotype-specific patterns and broader insights into variations between different groups. The overall response to vaccination within each study cohort was assessed by using the group-level comparison (**Figure 2**). The Shannon entropy index (SEI) was used to express transcriptomic diversity of modular repertoire functions at each contrast [16]. We first profiled the transcriptomic response to COVID-19 booster dose in the C-FV cohort by contrasting module abundance on days 1 and 5 against the baseline on day 0, observing a response involving 21 and 9 modules, respectively. On day 1, we observed a pronounced increase in six modules linked to interferon response and one module related to inflammation (M10.1, M15.86, M13.17, M15.64, M15.127, and M8.3 aggregate A28 and M13.12, aggregate A35 respectively, SEI = 0.79). Simultaneously, we noted a transient decrease in several modules associated with cytotoxic lymphocytes (M9.1 and M13.21), T cells (M15.38), cell cycle (M14.31, M15.73), and gene transcription (M14.17 and M14.69; SEI = 1.37), aligning with the established physiologic response to the BNT162b1 vaccine [17]. By day 5, we observed a rise in 9 modules, including transcripts associated with effectors of adaptive immunity such as plasma cells (M12.15, aggregate A27) and T cells (M12.6, M15.98, and M15.38, aggregate A1 and A6; SEI = 1.21). The comparison between day 5 and day 1 showed a polyfunctional response involving 92 modules, with a widespread upregulation of aggregates A1, A2, A3, and A5 (SEI = 2.19). These aggregates are associated with effectors of adaptive immunity and cell proliferation or activation, such as cytotoxic lymphocytes (M9.1 and M13.21), lymphocytes (M13.27, M16.78, M13.18, M15.24), T cells (M14.42, M15.38, M12.6, M15.98, M16.3, M15.33), cell cycle (M14.31, M12.3, M15.73), cell death (M15.29, M15.23, M15.49), gene transcription (M13.2, M14.71, M14.4, M14.34, M14.17, M14.23, M16.36, M15.21, M15.8, M15.3, M13.13, M14.69), mitochondrial stress/proteasome (M15.79), oxidative phosphorylation (M14.63), protein modification (M16.50, M16.99, M15.92, M16.13, M15.44, M12.1, M15.31, M12.5) and protein synthesis (M14.75, M11.1, M15.51, M13.4, M16.73, M10.2, M16.68, M16.56, M14.44). In concomitance with such widespread activation of immune response and cell machinery, modules associated with interferon and inflammation were downregulated (SEI = 1.76; **Figure 2**). Response values for each module at each time point contrast are reported in **Supplementary Table 5**.

**Figure 2.**
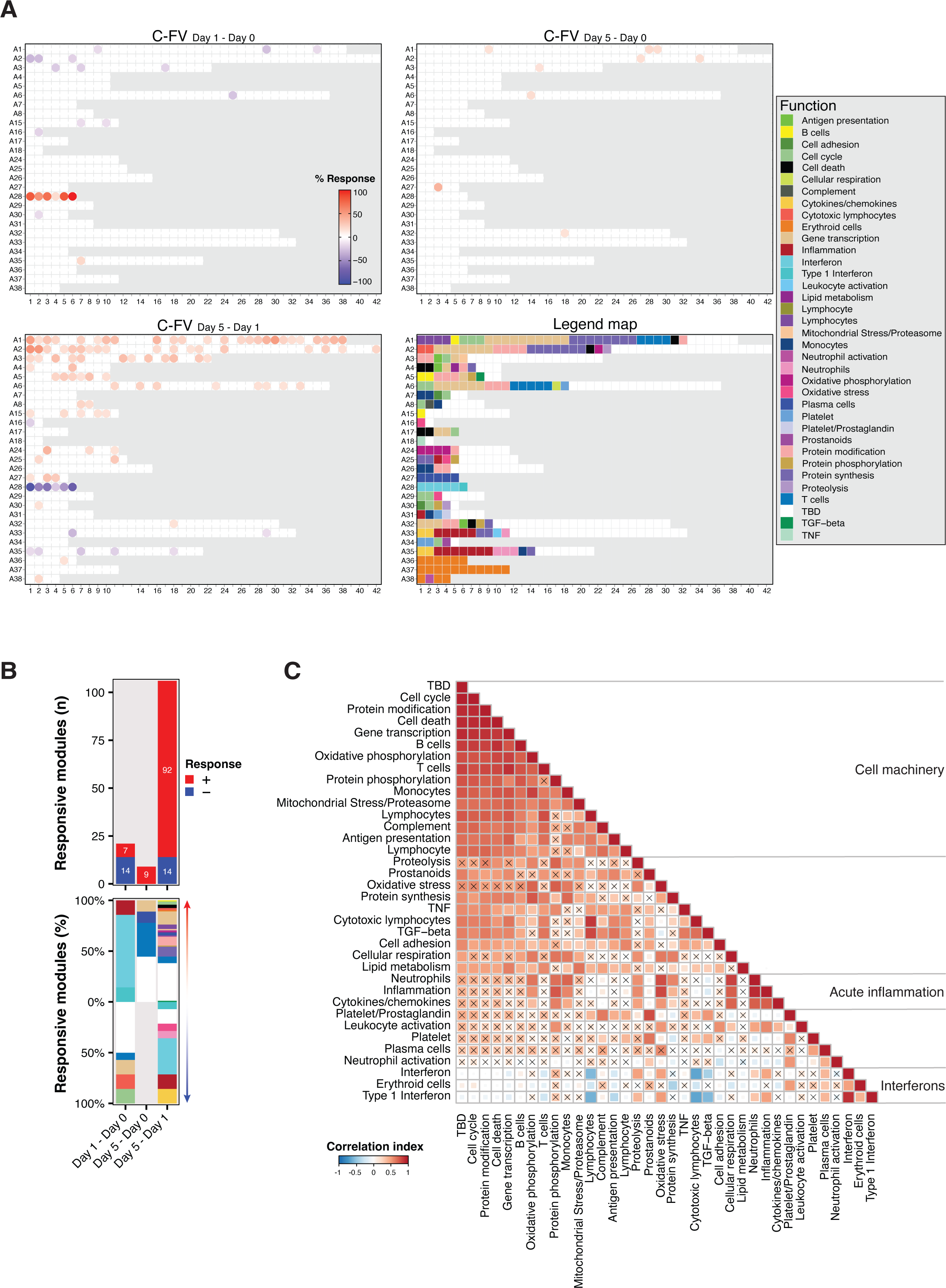
Transcriptional profile following the booster dose of the COVID-19 vaccine in the C-FV cohort. (A) Fingerprint plot displaying the BloodGen3 module responses on days 1 and 5 relative to day 0, and on day 5 relative to day 1 in the C-FC group. Modules are arranged in fixed positions, with each row (A1 to A38) representing a module aggregate. Red indicates modules with increased transcript levels compared to the reference time point; blue indicates decreased levels. The response percentage indicates the proportion of genes with increased or decreased abundance. Modules with a response between −15% and 15% are shown in white. (B) Bar plots summarizing the modular responses at each time point, with the upper plot showing the count of upregulated (red) and downregulated (blue) modules, and the lower plot showing the percentage of responsive modules. Each bar is color-coded by the functional category of the modules as indicated in the panel A legend. (C) Correlation matrix presenting Pearson correlation coefficients among BloodGen3 functional signatures, calculated using day 1 responses at the individual sample level.

To further explore the functional trajectories of COVID-19 booster, modules sharing the same functional annotation in Bloodgen3 were incorporated and assessed in a correlation matrix (**Figure 2C**). Notably, interferon modules had a significant positive correlation with plasma cells and inflammation and were inversely correlated with cytotoxic lymphocytes, lymphocytes, and TGF-beta. Signatures associated with cell machinery processes (i.e. cell cycle, cell death, gene transcription, oxidative phosphorylation, mitochondrial stress/proteasome, protein modification and phosphorylation) clustered with signatures of either innate (i.e. monocytes, complement, antigen presentation) or adaptive (i.e. B cells, T cells, lymphocytes) immunity. In parallel, we observed a further cluster of signatures associated with acute inflammation (i.e. neutrophils, inflammation, cytokines/chemokines).

Taken together, these findings indicate that the COVID-19 booster vaccination triggers an immediate, transient interferon surge, driving an orchestrated, polyfunctional adjustment of adaptive and innate immunity pathways.

### Comparative transcriptomic responses to COVID-19 booster reveal distinct patterns and functional dynamics in ICI and RTX patients

We assessed the transcriptomic response to COVID-19 booster in ICI and RTX patients (**Figure 3**; **Supplementary Figure 4**). The magnitude of modular response, in terms of absolute number of upregulated or downregulated modules, and the transcriptomic diversity of modular repertoire functions, in terms of SEI, were compared with C-FV at each time point contrast through Fisher’s exact test and Hutcheson test, respectively.

**Figure 3.**
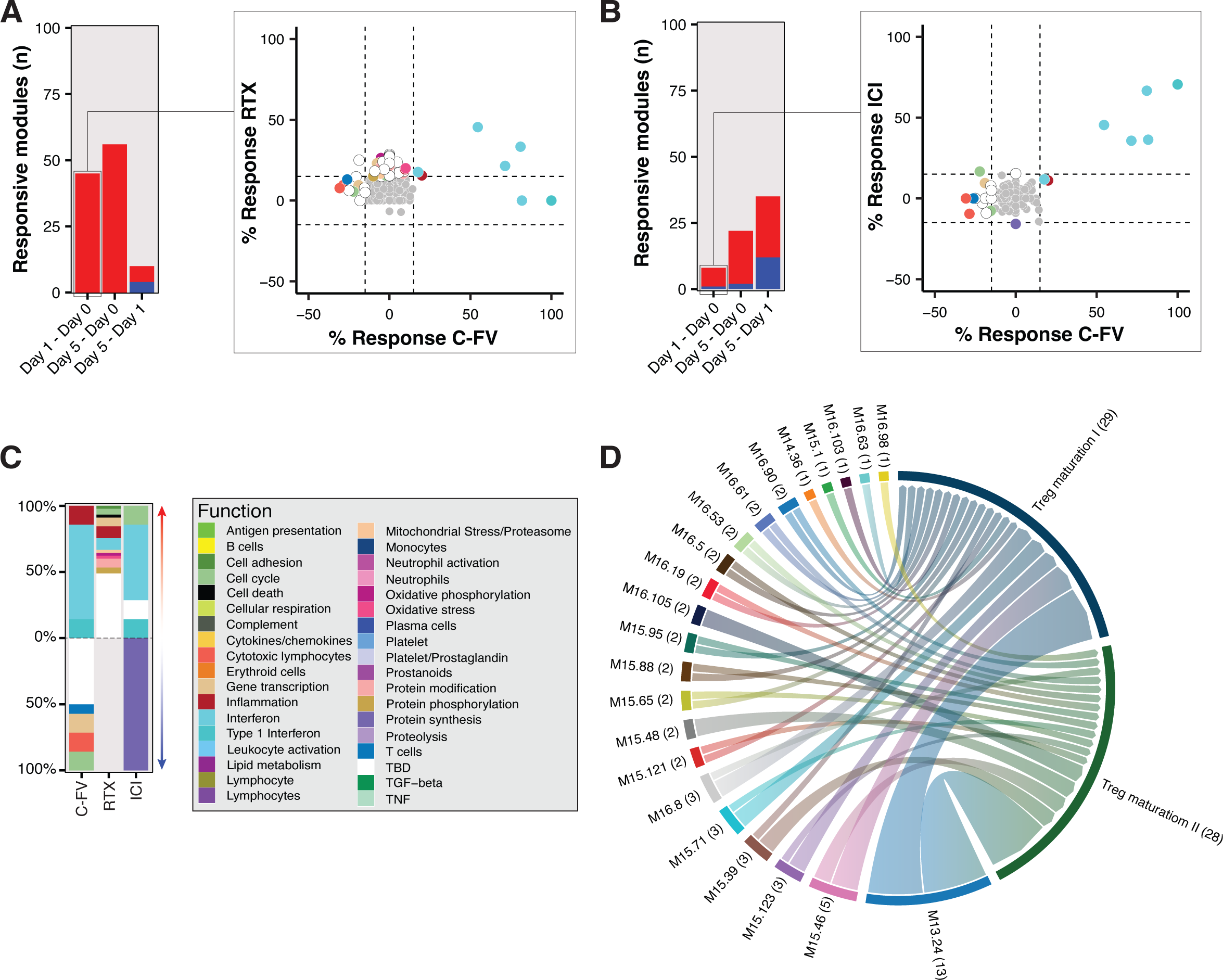
Comparison of transcriptomic responses to the COVID-19 booster in C-FV and cancer patients. A), B) Bar plots displaying the number of upregulated (red) and downregulated (blue) modules at each time point contrast for RTX (A) and ICI (B) patients. The scatter plots show the percentage of module response on day 1 relative to day 0 for RTX and ICI patients (y-axis) against C-FV (x-axis). Each point, representing a Bloodgen3 module, follows the color scheme from Figure 2 for functional annotation, with points colored only if the response exceeds 15% or falls below −15%. C) Bar plots illustrating the relative module response for C-FV, RTX, and ICI patients on day 1 compared to day 0. D) Circos plot visualizing gene overlaps between upregulated TBD modules in RTX patients on day 1 and specific signatures, and highlighting the interconnectedness of responses across different patient groups.

After booster administration, ICI patients showed a day 1-transcriptomic response similar to C-FV individuals with upregulation in 8 modules (p-value = 1) and a predominant spike of interferons (SEI = 1.15, p-value = 0.18; **Figure 3**) while having, in contrast to C-FV, only one downregulated module (p-value = 0.0008), associated with protein synthesis (M13.28). On day 5, ICI patients displayed upregulation in 20 modules (p-value = 0.057) including transcripts linked to cell cycle (M14.64 and M14.31), plasma cells (M15.110, and M15.12), platelets (M14.81, and M15.45), monocytes (M15.7), T cells (M12.6), and gene transcription (M14.55; SEI = 1.95, p-value 0.005; **Supplementary Figure 5A**). Comparative analysis of module abundance between day 5 and day 1 revealed a response of 23 modules (p-value = 1.51 10^-12^) involving functions related to the adaptive immunity and cell proliferation, including cytotoxic lymphocytes (M13.21), cell cycle (M14.64, M16.104), protein synthesis and modification (M14.58, M13.28, M14.13), T cells (M15.34), and plasma cells (M12.15; SEI = 1.93, p-value = 0.14; **Supplementary Figure 5B**). Modules associated with interferon and inflammation were found deregulated (SEI = 1.54, p-value = 0.19), aligning with C-FV.

RTX patients exhibited a distinct early immune perturbation, with the upregulation of 45 modules occurring at day 1 (p-value = 2.58 10^-8^), including transcripts associated with oxidative phosphorylation (M14.13), gene transcription (M14.34, M15.126, M14.67), cell adhesion (M15.56), protein modification (M15.92,M15.44, M14.14) and phosphorylation (M13.19, M16.84), oxidative stress (M16.107), and inflammation (M15.37, M14.19, M13.12, and M14.24; SEI = 1.84, p-value = 0.003; **Figure 3**). Notably, 22 of the 45 upregulated modules were not functionally annotated in Bloodgen3 (to be determined - TBD). We therefore performed overrepresentation analysis of transcripts included in such modules by using MSigDB [18,19], observing a significant association with two signatures related to the differentiation of T cells towards regulatory T cells (GSE42021_TREG_VS_TCONV_PLN_UP, FDR = 0.047; GSE42021_CD24LO_TREG_VS_CD24LO_TCONV_THYMUS_DN, FDR = 0.047; **Figure 3D**). On day 5, there was an upregulation of 56 modules (p-value = 4.31 10^-10^) pertaining to both adaptive and innate immunity, including T cells (M15.98), neutrophils (M14.28), inflammation (M14.19, M14.50, and M14.24), oxidative stress (M16.107), and B cells (M16.95), along with modules associated with cell proliferation and death (M14.54 and M16.97) and protein machinery (M14.14, M15.44, M12.1, M16.84, M13.10, and M16.11; SEI = 1.70, p-value = 0.03; **Supplementary Figure 5A**). Thirty-three of 56 modules were TBD, but no significant associations with MSigDB signatures were observed. The differential module response assessment at day 5 vs. day 1 highlighted a positive response involving only 6 modules (p-value = 2.88 10^-23^), with the upregulation of two modules linked to platelets (M16.109 and M10.3), two to prostaglandins and prostanoids (M16.64, M15.102), and one to the cell cycle (M14.39; SEI = 1.56, p-value = 0.001; **Supplementary Figure 5B**). Four modules, including two associated with interferon (M10.1, M15.86) and one associated with cell cycle (M13.20), were found deregulated (SEI = 1.04, p-value = 0.005).

Overall, ICI’s response aligned with that of C-FV, with a more pronounced day 5-response in terms of both magnitude and diversity. Conversely, RTX patients exhibited a markedly divergent phenotype of response from C-FV, as highlighted by the transcriptional entropy occurring on both day 1 and day 5 compared to day 0 and the modest response occurring between day 5 and day 1, denoting an uncoordinated response to vaccination.

### Type I interferon response to COVID-19 booster is blunted in RTX patients

Despite the different response patterns and treatment conditions, all study cohorts exhibited a notable interferon peak on day 1 post vaccination. Therefore, we set to profile in greater detail the response dynamics of aggregate A28 across the different groups (**Figure 4**). To this aim, we used two distinct signatures identified in our previous work by performing hierarchical clustering on the six interferon modules of aggregate A28 after the first and second dose of COVID-19 vaccine [11]. Of these, signature A28/S1 includes modules M8.3, M10.1, and M15.127, while signature A28/S2 includes modules M15.64, M13.17, and M15.86.

**Figure 4.**
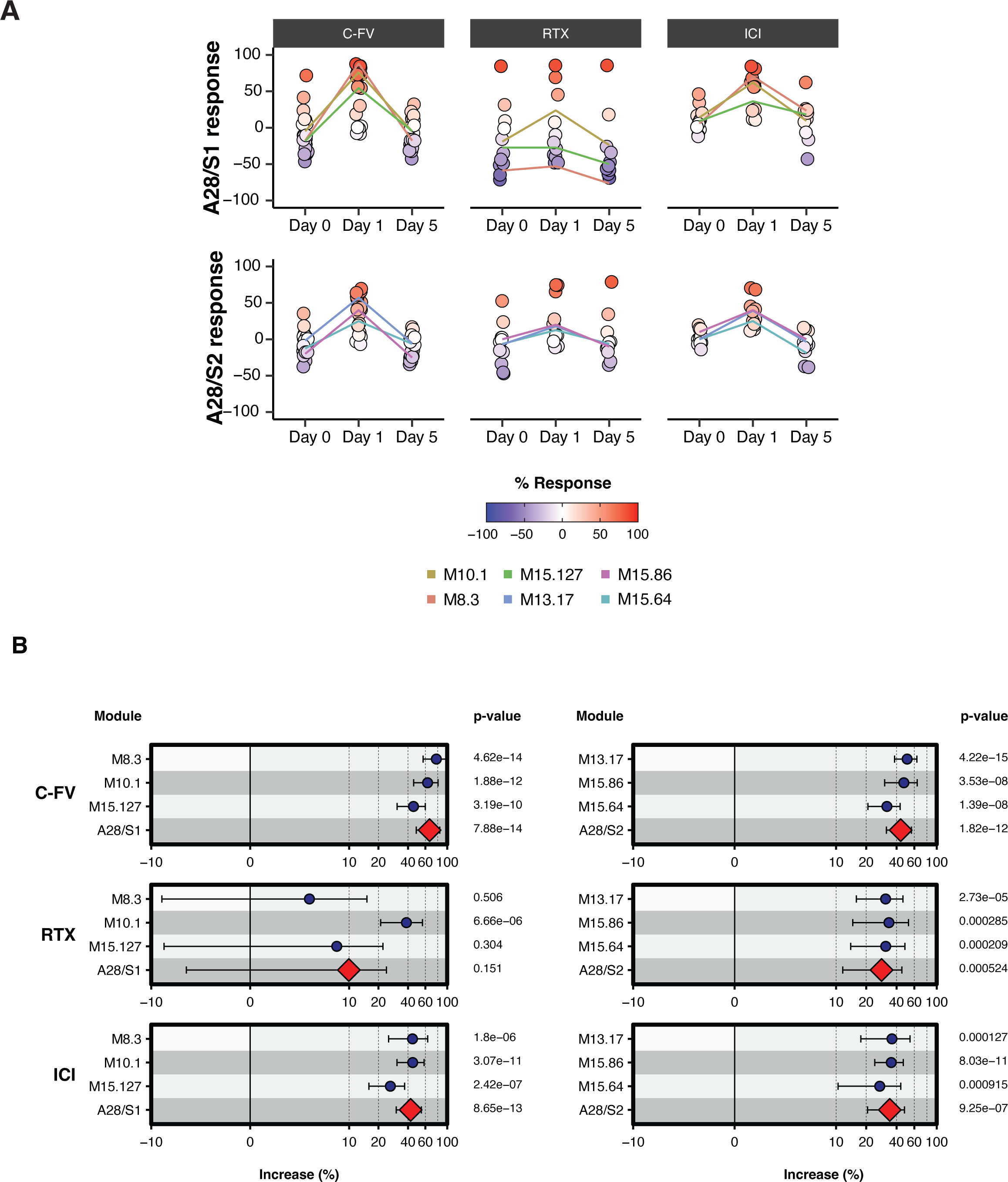
Dissection of the interferon response in the different study cohorts. A) Dot plots showing the responses of A28/S1 (upper panel) and A28/S2 (lower panel) across study cohorts, with individual sample-level assessments. Each dot’s color corresponds to the response percentage of A28/S1 or A28/S2, and lines indicate the average response for each module at each time point. B) Forest plot illustrating the interferon response on day 1 across cohorts, detailing the effects on A28/S1 and A28/S2 signatures and their modules.

By implementing a mixed effect model to account for the response of each signature between day 1 and day 0, we identified A28/S1 as significantly increased in C-FV (p-value = 7.88 10^-14^) and ICI patients (p-value = 8.65 10^-13^). Notably, RTX patients did not mount a significant response in the A28/S1 signature (p-value = 0.151). In particular, of the A28/S1 modules, the response of M8.3 (p-value = 0.506) and M15.127 (p-value = 0.304) were depleted, while the response of M10.1 was preserved (p-value = 6.66 10^-6^). A28/S2, along with each module included in the signature, was found significantly increased in all cohorts.

A28/S1 modules encompass genes from the oligoadenylate synthetase family (*OAS1*, *OAS2*, *OAS3*, *OASL*), interferon-induced protein family (*IFI6*, *IFI27*, *IFI35*, *IFI44*, and *IFI44L*), and interferon-induced protein with tetratricopeptide repeats family (*IFIT1*, *IFIT3*, and *IFIT5*), canonically induced by type I interferon. Modules of signature A28/S2 include transcription factors *IRF9* and *STAT2*, along with nuclear antigen family members *SP100*, *SP110*, and *SP140*, associated with interferon-γ signaling pathway.

Such results indicate a substantial impairment of the type I interferon response in RTX patients, a key component in the innate response to viruses and the cross-talk between innate and adaptive immunity [20,21]. This impairment in the type I interferon signaling may underpin the uncoordinated response to the COVID-19 booster observed for such patients, resulting in a dysfunctional response to the vaccine with a switch of T cells towards a regulatory phenotype [22].

### Immune celI deconvolution of blood transcripts reveals depletion of antigen presenting cells and B cells in RTX patients after booster administration

In order to evaluate changes in blood cell composition after booster administration across the different study cohorts, we performed immune cell deconvolution on blood RNA-seq. To this aim, multiple deconvolution methods were assessed. CIBERSORT was identified as the method that showed the highest correlation with peripheral blood counts performed according to clinical practice concurrently with RNA-seq and was therefore chosen for further analysis (**Supplementary figure 6**).

In the C-FV cohort, we observed an increase of plasma cell and memory B cell on day 1 and day 5 compared to day 0 (**Figure 5, Supplementary figure 6**). Moreover, an initial increase in monocytes and activated dendritic cells was noted (**Figure 5, Supplementary figure 6**), in line with the available literature [10]. CD8^+^ T cells exhibited an upregulation on day 5 following a transient decrease on day 1.

**Figure 5.**
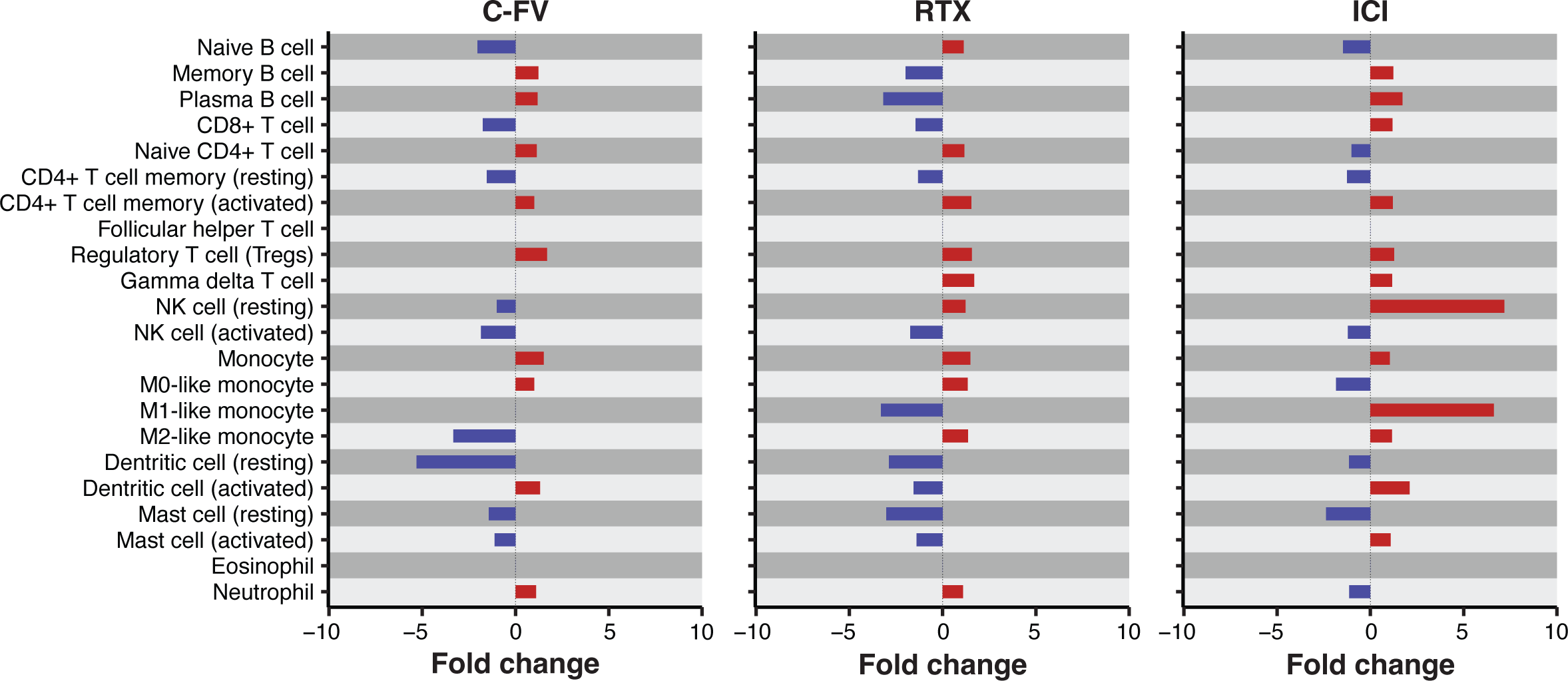
Changes in blood cell composition in the different study cohorts on day 1 compared to day 0. Bar plots illustrating the variation in immune cell populations, analyzed using CIBERSORT, on day 1 relative to day 0 for each study cohort. Each bar indicates the fold change of specific cell populations from the baseline, covering a range of immune cells including B cells, T cells, NK cells, monocytes, dendritic cells, and others, across C-FV, RTX, and ICI cohorts.

Similar patterns were noted in ICI patients, where blood cell deconvolution revealed a markedly pronounced upregulation of resting NK cells and M1-differentiating monocytes on day 1.

Of special interest, RTX patients displayed a suppression of antigen presenting cells, such as activated dendritic cells and M1-differentiating monocytes, as well as B cells on both day 1 and day 5. Across all study cohorts, a progressive upregulation of CD4^+^ T activated memory cells was observed.

## Discussion

In the present study, we conduct the first comprehensive characterization of the early and late phases of the immune response to consistently reproducible antigenic stimulation in patients treated with anti-CD20 agents and ICIs, thus bridging a knowledge gap regarding the biological determinants that underpin their immunosuppressive and immunostimulatory effects. Specifically, we used the COVID-19 booster as a model to assess the immune response in two distinct cohorts of patients with blood and solid cancers undergoing active treatment with rituximab and ICIs. To this aim, we opted for a system-scale approach based on the analysis of co-dependent transcriptional networks identified by Altman *et al*. [14]. Differently from traditional methods based on feature selection or dimension reduction, this approach considers the complex interdependence of variables in biological systems, allowing for a more informative evaluation of transcriptional changes that would not be detected if considering single genes as independent variables [23].

Our initial step was to characterize the immune response to COVID-19 vaccination in individuals with good overall health (C-FV). The booster dose of vaccine in the C-FV cohort led to an early isolated interferon response resembling the one observed after the first dose [11]. This finding confirms that the temporal proximity of the two vaccine doses administered within the primary cycle is critical to elicit the polyfunctional innate response seen after the second dose [11], even in individuals who have completed the primary vaccination cycle. The early isolated interferon peak following the booster dose of vaccine likely acts as a trigger of immune memory mechanisms involving cells of the adaptive immunity primed by the first and second doses [24]. Such transcriptional response is then associated with the restoration of SARS-CoV-2 humoral immunity, as highlighted by the marked increase of vaccine-targeted antibodies after four weeks.

Similarly to C-FV, ICI patients showed a preserved humoral and cell-mediated immune response to COVID-19 booster, while RTX patients were heavily immunosuppressed. Blood transcriptome dynamics in ICI patients resembled that of C-FV, with subtle differences consistent with a state of ongoing immune stimulation, including the absent downregulation of immune modules on day 1, observed for the C-FV, and an enhanced response on day 5. The marked upregulation of NK cells observed by cell deconvolution on day 1 aligns with data reported by Nelli *et al*. [25] and, together with the pronounced response of M1-differentiating monocytes, further supports such hypothesis. No other unusual findings were observed in the dynamics of humoral, cellular and transcriptomic response of ICI patients, supporting the efficacy and, likely, the safety of the concomitant administration of immunotherapy and COVID-19 vaccines.

On the other hand, the transcriptional profile of RTX patients was characterized by an overall immune dysfunction, denoted by the state of transcriptomic perturbation observed on both days 1 and 5, coupled with a substantial deficit in the type I interferon response and the upregulation of transcripts associated with differentiation of regulatory T cells. Type I interferon plays a critical role in the innate response to viruses, including SARS-CoV-2 [20], and is implicated in the crosstalk between antigen presenting cells and B cells, enhancing the humoral immunity by promoting B cell differentiation and isotype switching [21]. Consistently, immune cell deconvolution of blood transcripts showed a concomitant depletion of antigen presenting cells and B cells in RTX patients. Notably, Graalmann *et al*. described a type I interferon-dependent mechanism in the impairment of CD8^+^ T cell responses following B-cell depletion in murine models infected with modified vaccinia virus Ankara [26]. In parallel, the administration of rituximab has been associated with expansion of Tregs, likely as a direct effect of B cell depletion [7]. Of interest, depletion of Tregs has been associated with an enhanced response to COVID-19 vaccination, suggesting an inverse correlation with vaccine efficacy [27].

Notably, the humoral and cell-mediated responses assessed six months after the booster dose may have been influenced by intercurrent immune stimulations. Overall, the COVID-19 booster effectively enhances humoral response against Spike protein antigens, with no observed effect on anti-Env and anti-Nucleo antibodies, aligning with existing evidence [11]. Vaccine-targeted antibodies remain persistently elevated six months post-booster. While this suggests that the booster dose elicits long-lasting humoral immunity to the virus, the significant rise in infection-associated antibodies at month 6 implies a role of natural infection in stimulating and maintaining a polyfunctional humoral response to the virus. Indeed, seroconversion for Env and Nucleo antigens in most C-FV at month 6, irrespective of COVID-19 diagnosis, likely reflects asymptomatic COVID-19 cases or, intriguingly, infection with cross-reactive coronaviruses, as indicated by the significant positive correlation of SARS-CoV-2 IgMs with anti-S1 IgMs of hCoVs six months after vaccination. In such regard, while it is unclear whether pre-existing hCoVs immunoglobulins confer protection towards COVID-19, cross-reaction between SARS-CoV-2 and hCoVs has been documented [28,29]. Since SARS-CoV-2 infection results in increased titers of hCoVs antibodies [30], infections by common hCoVs may in turn enhance the humoral immunity towards SARS-CoV-2, providing indirect protection by acting as a natural booster of the host immune response.

Our work has several limitations. The small size of our sample highlights the exploratory nature of our study, particularly in assessing cell-mediated immunity. This limitation restricted our ability to evaluate and adjust for possible confounding factors like age and comorbidities. In our cohort, cancer patients were generally older and had more comorbidities than those in the C-FV group. Despite recognizing this limitation, the rapid deployment of the booster dose in Italy necessitated swift action within a very limited timeframe, which explains our lower participant numbers. Additionally, our assessment of humoral and cell-mediated immunity focused on the original SARS-CoV-2 variant without accounting for the emergence of new variants of concern. Nevertheless, this should not detract from the significance of our findings. Our goal was to assess the framework of immune response following a reliable, reproducible immune stimulus in patients treated with specific immune perturbagens. As such, our work may in principle be confirmed by downstream studies conducted during future vaccination campaigns. Finally, blood cell decomposition was performed computationally on blood RNA and can therefore reflect changes in the transcriptional activity of cells rather than in the actual blood composition. Nonetheless, changes in blood cell deconvolution values can be indicative of underlying changes in cell function or state and may still be relevant to understanding the response to various conditions or treatments in different groups. Indeed, the increase of NK cell and M1-differentiating monocytes in ICI patients and the decrease of antigen presenting cells in rituximab patients offer intriguing and biologically coherent insights into the interplay amongst the various actors of innate and adaptive immunity and its alterations in non-physiological conditions.

In conclusion, our study represents a significant advancement in understanding the immune responses to COVID-19 vaccination in patients undergoing treatment with immune-perturbing agents such as RTX and ICIs. By employing a system-scale analysis of transcriptional networks, we were able to uncover intricate details of the immune response that traditional methods might overlook. Our findings reveal that the early interferon response plays a pivotal cross-talk role in triggering immune memory mechanisms, crucial for the restoration of SARS-CoV-2 humoral immunity. The preserved humoral and cell-mediated responses in ICI patients, contrasted with the immune dysfunction in RTX patients, highlight the potential impact of these treatments on vaccine efficacy. Moreover, our work points to the importance of type I interferon responses and the regulatory role of T cells in shaping the B-mediated immune response to vaccination. Likewise, our findings stress the relevance of a functional B-mediated response to elicit an orchestrated innate and adoptive T-cell mediated immunity. Our efforts lay the groundwork for future studies aimed at optimizing vaccination strategies and improving outcomes in vulnerable populations, as well as understanding better how B and T cells interact in coordinating the immune response in physiological and pathological states. Further research is needed to validate our results in larger cohorts and against emerging variants of concern, but our study marks a crucial step towards understanding the biological determinants of immunosuppressive and immunostimulatory effects of cancer therapies on vaccine-induced immunity.

## Methods

### Ethics statement

The present work has been reviewed and approved by the ethics committees of Istituto Nazionale per le Malattie Infettive Lazzaro Spallanzani (INMI, Rome, Italy) and Regione Liguria (Genoa, Italy; ID: 12131). Prior to study participation all recruited patients and volunteers provided written informed consent. All procedures carried out in the present work are in accordance with the 1964 Helsinki Declaration and its later amendments.

### Patient recruitment

Adult patients with solid or blood cancer were recruited at the Hematology, Oncology, and Internal Medicine units of IRCCS Ospedale Policlinico San Martino (HSM - Genoa, Italy). C-FV were recruited amongst health care workers at University of Genoa and HSM, as well as external volunteers in good general health conditions. All subjects were scheduled to receive the third dose of COVID-19 mRNA vaccine, either BNT162b2 or mRNA-1273. RTX patients were considered eligible for the study if treated with rituximab within the previous nine months. ICI patients were considered eligible if undergoing active treatment with ICI at the moment of recruitment.

All subjects were assessed using the following exclusion criteria: symptomatic infection of proven or suspect viral or bacterial origin in the previous 8 weeks (paraneoplastic fever was not considered an exclusion criteria); major trauma or surgery in the previous 8 weeks; hereditary immunodeficiency; HIV infection; active immunologic disorders (autoimmune disease, active allergic disease or undergoing chronic treatment); treatment with immunosuppressive drugs not for cancer treatment; acute or chronic end-stage liver disease; acute or chronic end-stage (stage III-IV) kidney disease.

### Study design

Capillary blood samples were collected from all subjects at day 0, day 1, day 5, week 4 and month 6 after vaccination. Peripheral venous blood samples were collected from a subset of patients and C-FV at day 0, day 5, week 4, and month 6. No patient received cancer treatment between day 0 and day 5. Whole blood transcriptome sequencing was performed on capillary blood samples collected at day 0, day 1, and day 5. Multiplex SARS-CoV-2 and hCoV serology was performed on capillary blood samples collected at day 0, week 4, and month 6. Cell-mediated immunity to SARS-CoV-2 was assessed through interferon-γ release assay (IGRA) on peripheral venous blood samples collected at day 0, day 5, week 4, and month 6. Intercurrent diagnoses of COVID-19 were assessed at each appointment scheduled per study protocol. Subjects were considered as diagnosed with COVID-19 in case of positive molecular or antigenic nasopharyngeal swab recorded in HSM or reported at the moment of sample collection.

### Sample collection

For whole blood RNA-seq, we collected 50 µL of capillary blood through finger-puncture using the capillary blood and tube assembly GK100 (Kabe Labortechnik, Nümbrecht, Germany), previously aliquoted with 100 µL of Tempus™ Blood RNA Tubes stabilizing reagent [31]. GK100 tubes were then labeled and stored at −20°C in a dedicated freezer (Beko). For multiplex SARS-CoV-2 and HCoV serology, we collected 20 µL of capillary blood through finger-puncture using Mitra® fingersticks (Neoteryx, Torrance, CA, USA), subsequently stored at room temperature. Peripheral venous blood samples (∼5 mL) for IGRA were collected through peripheral venipuncture in one heparinized Vacuette ® tube.

### Whole blood RNA extraction, quality check, and library preparation

Total RNA was extracted from GK100 tubes using the Tempus^TM^ Spin RNA Isolation Kit Protocol with minor modifications. RNA quality check was performed using the RNA ScreenTape (Agilent) on a TapeStation 2200 device (Agilent). RNA samples with RNA integrity number above 7 were considered of good quality. Up to 200 ng of input RNA were used to prepare the libraries for 3’ mRNA-Seq using the QuantSeq 3’ mRNA-Seq Library Prep Kit for Illumina FWD with UDI 12 nt Unique Dual Indexing (Lexogen). Libraries were outsourced to Lexogen for sequencing at 1×75 bp length, 8 megareads per sample on a NextSeq1000 sequencer.

### Assessment of the humoral response to SARS-CoV-2 and hCoVs

A custom bead array was developed to detect antibodies against specific human coronavirus proteins in serum samples. This array utilized carboxymethylated bead sets with varying intensities of an ultraviolet-excitable dye. Each bead set was conjugated to three distinct SARS-CoV-2 proteins: the envelope protein, the nucleoprotein, and the full-length spike protein or its fragments, including the RBD, the S-Trimer, and S1. Additionally, the S1 subunit of closely related viruses including SARS-CoV, MERS, HCoV-229E, HCoV-NL63, HCoV-HUK1, and HCoV-OC43 was evaluated. The binding of human antibodies to each viral antigen (bead set) was visualized using fluorescently labeled isotype-specific mouse monoclonal or polyclonal antibodies. The levels of total IgM, total IgG, and total IgA as well as their individual isotypes IgG1, IgG2, IgG3, IgA1, and IgA2 were measured. The assays were conducted on filter plates and analyzed using a BD Symphony A5 instrument with a high-throughput sampler. An average of 300 beads per region was analyzed, and the median fluorescence intensity (MFI) for each isotype binding was employed to characterize the antibody response. A binding antibody index was calculated by dividing the MFI of pooled negative blood controls collected before June 2018 (Sidra IRB 1609004823) by the MFI obtained for vaccinated donor samples.

### Assessment of cell-mediated immune response to vaccination

The cell-mediated immunity to SARS-CoV-2 was evaluated through IGRA using a commercially available kit (Quan-T-cell Euroimmun^TM^). Fresh whole blood samples were aliquoted in three different tubes respectively with a) no T-cell stimulation for the assessment of the interferon-γ background; b) specific T-cell stimulation using SARS-CoV-2 S1; c) nonspecific T-cell stimulation by means of a mitogen. All samples were stimulated within five hours from collection as per manufacturer’s instructions. After the incubation, each tube was centrifuged to obtain stimulated heparinized plasma. The concentration of interferon-γ was subsequently assessed using the enzyme-linked immunosorbent assay (ELISA) provided by the manufacturer. We used the recommended thresholds of interferon-γ concentration to determine negative, borderline, and positive results, respectively set at < 100 mUI/mL, 100-200 mUI/mL, > 200 mUI/mL.

## Computational methods

### Assessment of binding antibody index changes

The matrix of raw antibody indices was processed within the R environment using custom scripts. First, the fold-change for each time point against the relative baseline for each antibody was assessed at individual sample level. To estimate group variations compared to the basal level of each antibody, we implemented a mixed-effects model, accounting for paired samples and COVID-19 diagnosis intervening between samplings. The fixed model coefficients were extracted and used as the unitary increment of the antibody index as a function of time (*BAI_t_*) and COVID-19 diagnosis (*BAI_COVID-19_*). The correlation between SARS-CoV-2 and hCoVs antibodies was assessed with Spearman’s rank correlation coefficients using the raw antibody index matrix.

### Alignment of 3’ mRNA-Seq, quality check, and processing

FASTQ files were processed through the QuantSeq 3‘ mRNA-Seq Integrated Data Analysis Pipelines on the Bluebee® Genomics Platform. Using the QuantSeq FWD analysis pipeline, raw data were aligned by the STAR v2.5.2a to human genome GRCh38/hg38 (Ensembl release 77, August 2014). QC was performed into two steps, first on the raw counts using FASTQC for a preliminary report of sequencing results and trimming step, then using RSEQC for evaluation of distribution of the reads on the annotation features. Gene counts were estimated by HTSeq-count v0.6.0 and annotated using the Ensembl v77 GTF. All downstream analyses were performed in R. Raw counts were normalized by dividing for sample-specific size factors using the DESeq2 package.

### Immune modular repertoire analysis

Normalized transcriptional levels were aggregated using a prespecified set of 382 transcriptional modules as described in detail in [14,15], using the BloodGen3Module R package. In brief, based on the weighted clustering networks of co-expressed transcripts derived from 16 blood samples, a repertoire of immune modules was identified. Downstream analysis and visualization were performed using custom R scripts based on DESeq2, ggplot2, and ComplexHeatmap. The workflow started with annotating a normalized expression matrix with immune functionals modules, then differential expression was assessed and eventually the immune response was calculated. The immune response was quantified with two different approaches, namely “group comparison”, i.e. the proportion of constitutively expressed genes whose abundance levels differ significantly between two groups, or as “individual comparison”, defined as the proportion of constitutively expressed genes whose abundance levels differ significantly for the same individual compared to a reference point. The values range from −100% to 100%, according to the proportion of decreased or increased constitutive transcripts. The main steps for the group comparison are: a) estimate fold-change variation and p values from the mixed effect model for each gene, b) calculate number of genes belonging to the same module with positive or negative trends (|logFC| > 1.5, p value < 0.05), c) if the number of genes of the same module with a positive trend is greater than the number of genes with a negative trend, a positive response is assigned to the module, otherwise a negative response is attributed to the module. The response rate is calculated as the number of significant genes, considering the direction of the response, divided by the total number of genes of a given module. The main steps for the individual sample level comparison are: a) estimate fold-change of a given gene of a given sample considering as reference the average value of the same gene across a set of control samples, b) calculate the difference in terms of normalized counts between a given gene of a given sample against the average normalized counts of the same gene across a set of control samples, c) calculate number of genes belonging to the same module with positive or negative trends (|logFC| > 1.5, |absolute difference in counts| > 20), and d) if the number of genes of the same module with a positive trend is greater than the number of genes with a negative trend, a positive response is assigned to the module, otherwise a negative response is attributed to the module, in line with group comparison. By default, to enhance interpretability, only the strongest direction is displayed, with red shades depicting an increment whereas blue palette describing a decrease in the response. For the “group comparison”, we modified the BloodGen3 approach, implementing a linear mixed-effects model (lme4 package) that accounts for the paired nature of the design of the present study. The Blood3GenModule was also used to visualize the results for both group comparison and individual comparison analyses. As a quantitative measurement of the level of complexity of modular responses, we adopted the Shannon index, computed as the measure of diversity of module repertoire functions upregulated or downregulated at each time point contrast in the different study cohorts. SEIs were calculated using the ecolTest package and compared through multiple Hutcheson t-tests.

### Characterization of TBD modules in RTX patients

Characterization of genes associated to TBD modules upregulated in RTX patients were determined by over-representation analysis using the clusterProfiler package with default settings [32,33]. The C7 (version 7.4) immune signature collection from the MSigDB portal was adopted for annotation [18,19].

### Characterization of the day 1 interferon response

Modules of the aggregate A28 and signatures A28/S1 and A28/S2 were evaluated leveraging a mixed effect model, accounting for same-case sample pairing, measuring the temporal changes of response between day 1 and day 0 for each study cohort.

### Immune cell type in-silico deconvolution

To perform immune cell deconvolution on blood transcripts we assessed MCP counter, Epic, Quantiseq, Xcell, CIBERSORT, CIBERSORT absolute, and Abis, comparing neutrophil signatures assessed on blood RNA with paired neutrophil percentages in peripheral blood counts performed per clinical practice in a subset of 15 patients with same-day WBC and RNA-seq measurements. Neutrophils were selected as the benchmark for such comparison, being the immune subpopulation available for all deconvolution methods. The CIBERSORT algorithm was selected as the best performing one for downstream analyses (*immunedeconv* R package with defaults options).

## Declarations

### Data and code availability

Sequencing data, including raw counts and annotation, and raw antibody indexes are publicly available, together with custom R code, at Synapses.org (DOI: https://doi.org/10.7303/syn55333847). Further information and requests regarding wet laboratory experiments and data analysis should be directed to and will be fulfilled by the lead contact, Gabriele Zoppoli.

## Acknowledgments

The authors wish to thank all the patients and volunteers for making this study possible. This work was supported by Associazione Italiana per la Ricerca contro il Cancro (AIRC IG grant IG21570 to GZ), Associazione italiana contro le leucemie-linfomi e mieloma (AIL – to GZ and AB), Alleanza Contro il Cancro (ACC), CURIOSITY driven research funds (University of Genoa, to GZ), liberal donations, (Grazia Benvenuto e Piero Chiabra), Research Projects of National Interest (Italian Ministry of University and Research, ID:M4.C2.1.1 to GZ). The authors wish to thank Dr. P. Blandini, MD for his useful intellectual insights.

## Author contributions

Conceptualization: FR, NF, RML, DB, AB, GZ. Methodology: MD, GG, JCG, DB. Software DR, DC, LF. Validation: MD, LF. Formal Analysis: LF. Investigation: FR, MD, IL, MS, GG, NF, CS, BC, GG, AD, LZ, EC, GF, EM, AO, ABe, FF, BB. Resources: BB, JCG, DB, AB, GZ. Data Curation: FR, MS, BC, GF, LF. Writing – Original Draft: FR, MD, IL, MS, LF, GZ. Writing – Review & Editing: all authors. Visualization: FR, LF. Supervision: JCG, DB, AB, LF, GZ. Project Administration: LF, GZ. Funding Acquisition: AB, GZ.

## Declaration of interests

Carlo Genova declares honoraria from Astra Zeneca, Bristol Myers Squibb, Eli Lilly, Merck-Sharp-Dohme, Novartis, Roche, Sanof, Takeda. Davide Bedognetti is now employed at Kite Pharma (Los Angeles, CA, USA). Gabriele Zoppoli reports stock ownership of Immunomica Ltd.

## Supplementary material

### Supplementary Figures

**Supplementary Figure 1. Sample collection and event timelines across C-FV, RTX, and ICI cohorts.** This figure outlines the key milestones and sample collection points throughout the study period, spanning from 2020 through 2022. It includes timelines for vaccine administration (first dose, second dose, booster dose) and COVID-19 diagnosis events, alongside detailed sample collection moments (baseline, day 1, day 5, week 4, month 6). Each cohort’s timeline is distinctly marked, providing a clear visual representation of the study’s temporal framework and the critical points for data collection and analysis.

**Supplementary Figure 2. Humoral response to hCoVs in the different study cohorts.** This figure presents a comprehensive analysis of the humoral immune response against various hCoV antigens, including HCoV-229E-S1, HCoV-HKU1-S1, HCoV-NL63-S1, HCoV-OC43-S1, MERS-S1, and SARS-S1, across C-FV, RTX, and ICI cohorts. The response is measured in terms of log2 fold change in Binding Antibody Index (BAI) at weeks 4 and 6 months post-vaccination, with separate panels for each hCoV antigen. It includes data for IgM, total IgG, IgG1, IgG2, IgG3, IgA1, and IgA2, visually depicted to highlight differences in immune response magnitude and specificity within and between patient cohorts.

**Supplementary Figure 3. Correlations between SARS-CoV-2 and hCoV IgMs in the C-FV cohort.** The scatterplots depict the correlations between SARS-CoV-2 IgM targeting the Nucleo antigen (A), the RBD (B), the S1 (C), the ENV antigen (D), and the S-Trimer (E) and IgM targeting hCoVs S1.

**Supplementary Figure 4. Fingerprint grid graph depicting the transcriptomic response to the COVID-19 booster in RTX and ICI patients.** The figure is divided into panels comparing day 1 vs day 0 (A), day 5 vs day 0 (B), and day 5 vs day 1 (C) for RTX and ICI cohorts. Each panel represents a matrix with modules and response percentages, highlighting increases (red) and decreases (blue) in gene expression.

**Supplementary Figure 5. Comparative analysis of the day 5 transcriptomic responses in C-FV and cancer patients following the COVID-19 booster.** A) Comparison at day 5 versus day 0, showcasing the differential expression patterns and magnitude of response across the C-FV, RTX, and ICI cohorts. B) Day 5 versus day 1 comparisons, further delineating the temporal dynamics of gene expression changes induced by the booster shot.

**Supplementary Figure 6.** A) Correlation matrix of various immune deconvolution methods *versus* blood work results. B) Changes in blood cell composition by CIBERSORT deconvolution on day 5 compared to day 0 across the different study cohorts, using bar plots to display the fold change of the immune cell populations.

### Supplementary Tables

**Supplementary Table 1. Extended data of study participants.** Demographic and clinical information on the study participants across the different cohorts (C-FV, RTX, and ICI). Data points include participant ID, group classification, age, sex, tumor type (where applicable), histology, stage of disease, type of therapy received, and steroid assumption. Additionally, it includes information on steroid administration type, presence of autoimmune diseases, comorbidities, vaccination status, and specific dates related to the study timeline (e.g., vaccination and sample collection dates).

**Supplementary Table 2. Humoral response to SARS-CoV-2 in the different study cohorts.** This table details the statistical analysis of the humoral immune response to SARS-CoV-2, as measured by the Binding Antibody Index (BAI) across various immunoglobulins (IgM, IgG, IgG1, IgG2, IgG3) and their target antigen. For each antibody, the table provides the time regression coefficient, indicating the change in BAI over time, and the COVID-19 regression coefficient, reflecting differences in BAI due to COVID-19 diagnosis. Statistical significance is assessed through p-values for both BAI over time (BAIt) and BAI relative to COVID-19 (BAI COVID-19), with adjustments for multiple testing presented as False Discovery Rate (FDR) values. The “Time point contrast” column specifies the comparison timeframe, such as “Week 4 | Baseline,” facilitating an understanding of antibody dynamics post-vaccination.

**Supplementary Table 3. Humoral Response to human coronaviruses (hCoVs) in the different study Cohorts.** This table outlines the detailed analysis of the humoral immune response against antigens from various hCoVs, such as SARS-CoV, measured through the Binding Antibody Index (BAI) for different immunoglobulin classes (IgM, IgG, IgG1, IgG2, IgG3). It provides an overview of the time regression coefficient for each antibody, reflecting changes in BAI over the study period, alongside the COVID-19 regression coefficient, indicating the impact of a COVID-19 diagnosis on antibody levels. The statistical significance of these changes is evaluated by p-values for BAI over time and in relation to COVID-19 diagnosis, with corrections for multiple comparisons through False Discovery Rate (FDR) values. Each entry is contextualized within a specific time point contrast, such as “Week 4| Baseline”.

**Supplementary Table 4. Results of IGRA assays.** This table presents the outcomes of the Interferon-γ Release Assay (IGRA) for measuring immune responses in study participants across different cohorts (C-FV, RTX, ICI). It includes subject IDs, group categorization, and quantitative IGRA results (expressed in mIU/mL) at various time points: Day 0 (pre-booster), Day 5, Week 4, and Month 6 post-booster vaccination. Values exceeding the assay’s upper limit of quantification are indicated by “>1900 mIU/mL”.

**Supplementary Table 5. Immune module response at each time point contrast in the different study cohorts**. Responses of the immune modules, evaluated through transcriptomic analysis, across the time points contrasts (Day 1 vs. Day 0, Day 5 vs. Day 0, Day 5 vs. Day 1) in the study cohorts (C-FV, RTX, ICI). For each immune module, the table outlines the fold changes, statistical significance, and direction of response (upregulation or downregulation) at each contrasted time point.

## Notes

https://doi.org/10.7303/syn55333847

### Summary of Updates

The following version has been edited, including improved figures and supplementary material.

