## Supplementary figures and images for "Biological modifications of the immune response to COVID-19 vaccine in patients treated with rituximab and immune-checkpoint inhibitors"

### Supplementary figure 1

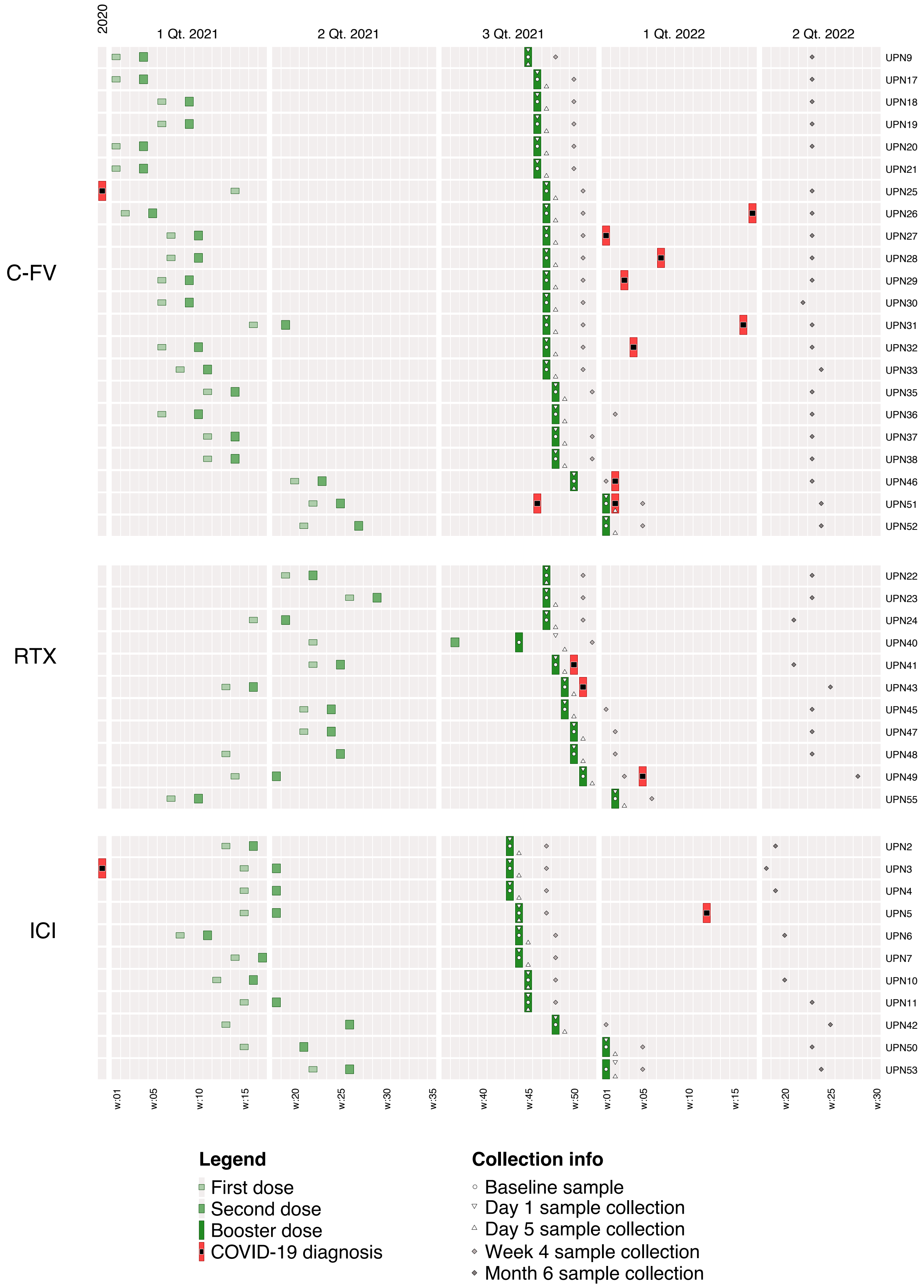

### Supplementary figure 2

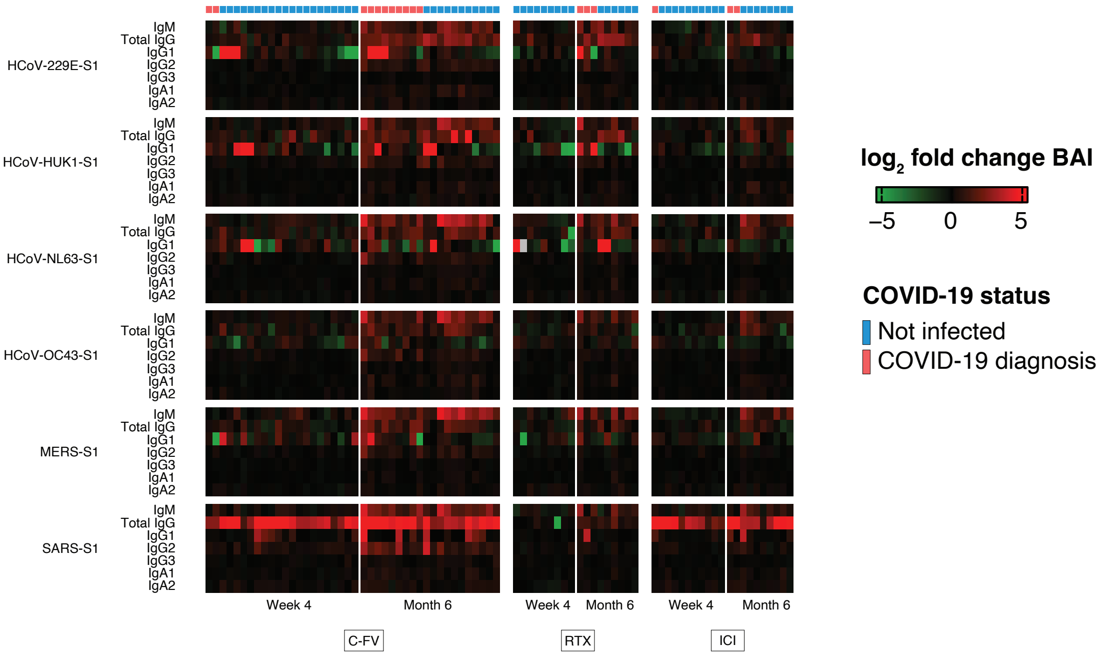

### Supplementary figure 3

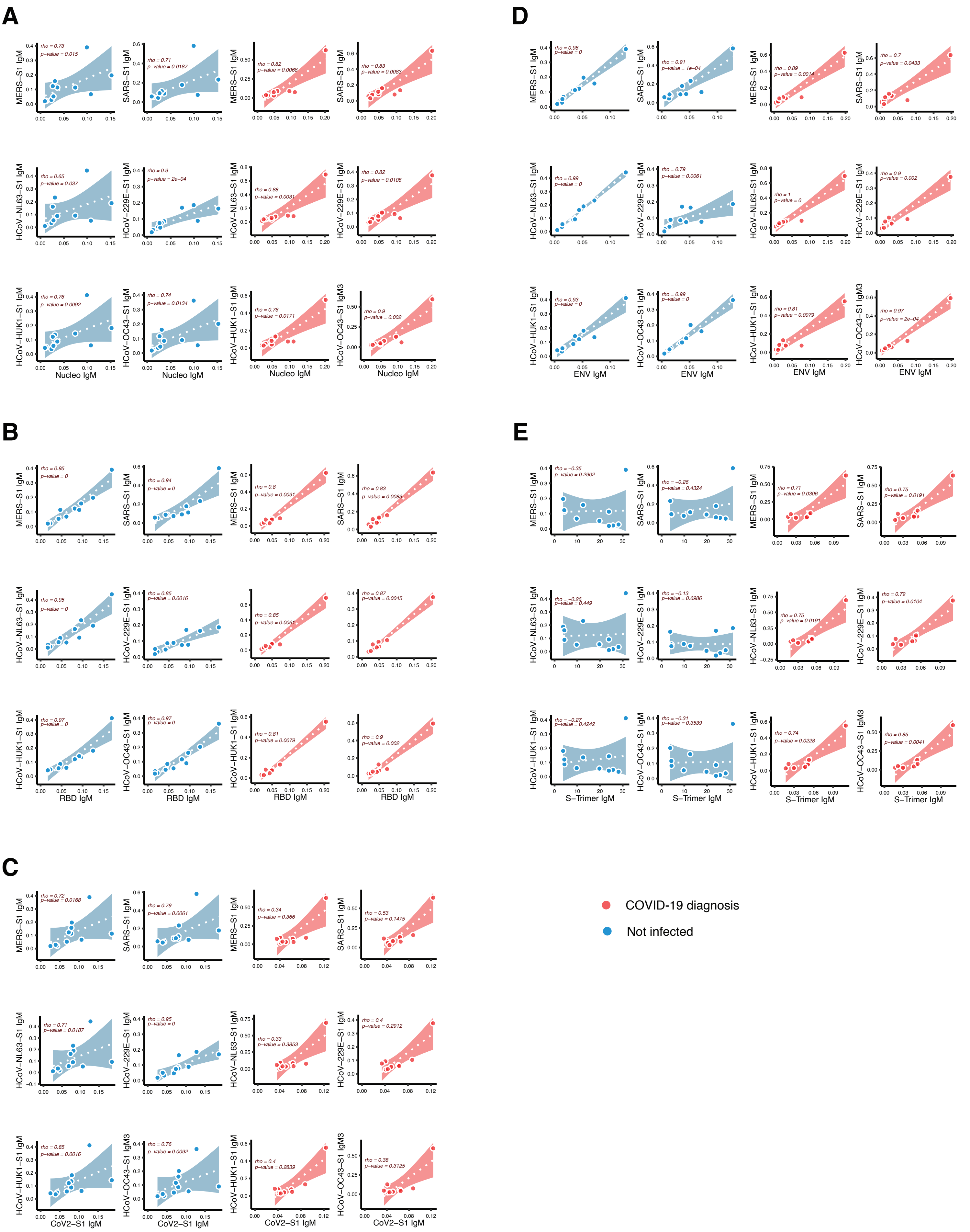

### Supplementary figure 4

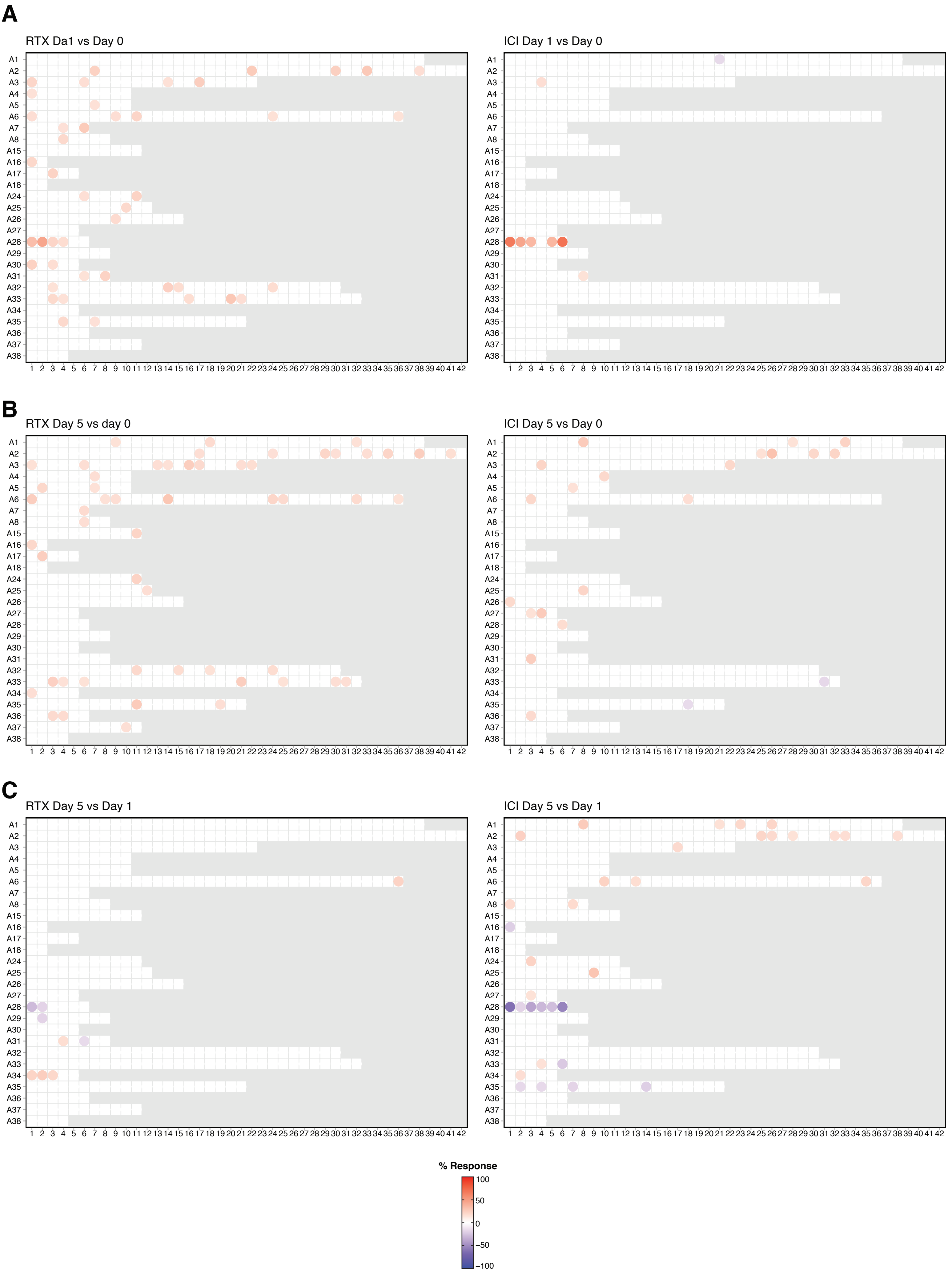

### Supplementary figure 5

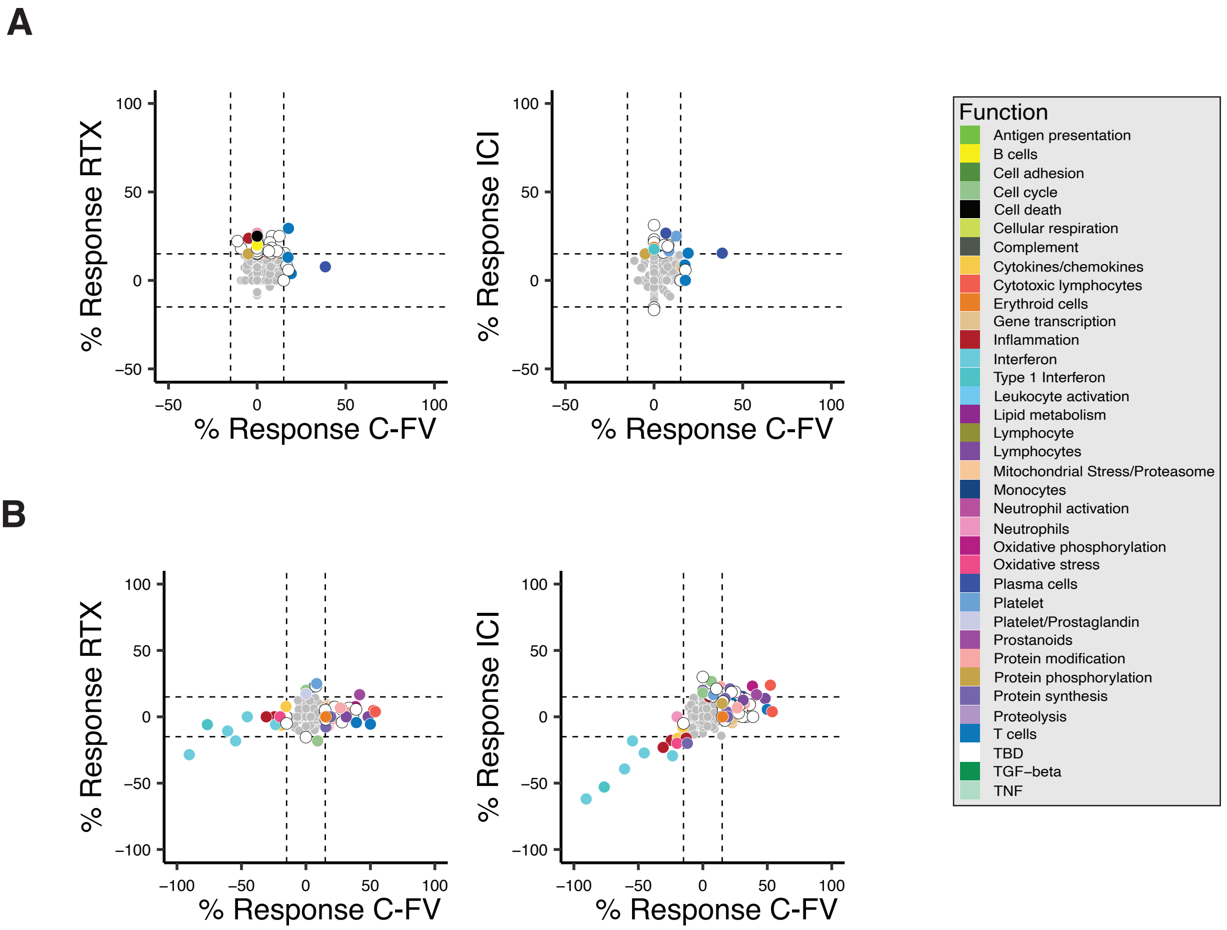

### Supplementary figure 6

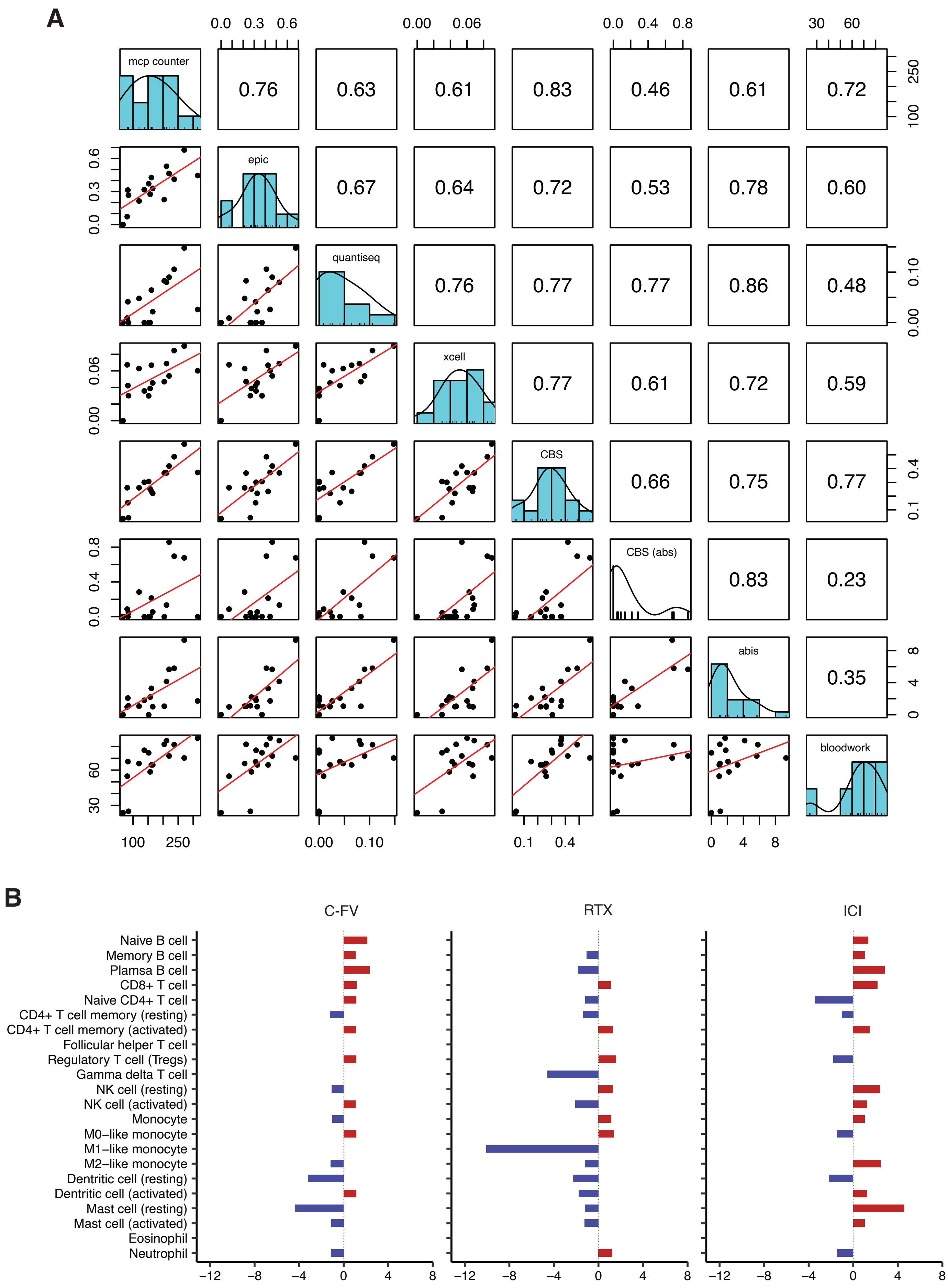
